# The representational geometry of out‐of‐distribution generalization in primary visual cortex and artificial neural networks

**DOI:** 10.1101/2025.04.23.650305

**Authors:** Zeyuan Ye, Ralf Wessel

## Abstract

Humans and other animals display a remarkable ability to generalize learned knowledge to novel domains, a phenomenon known as out-of-distribution (OOD) generalization. This capability is thought to depend on the format of neural population representations; however, the specific geometrical properties that support OOD generalization and the learning objectives that give rise to them remain poorly understood. Here, we examine the OOD generalization of neural population representations of static grating orientations in the mouse visual cortex. We show that a decoder trained on neural responses within a restricted orientation domain can generalize to held-out orientation domains. The quality of generalization correlates with both the dimensionality and the curvature of the underlying neural representation manifold. Notably, similar OOD-generalizable geometry emerges in a deep neural network trained to predict the next frame in natural video sequences. These findings reveal the representational geometric properties underlying OOD generalization, and suggest that predictive learning objectives offer a promising approach for acquiring generalizable representation geometry.

## Introduction

Animals, including humans, generalize knowledge across domains. For instance, a person who has learned to drive at low speeds can readily adapt those skills to high-speed driving on a highway. This capability for out-of-distribution (OOD) generalization is thought to rely on specific formats of neural representation^1–3^. While considerable progress has been made in understanding the neural representations underlying OOD generalization in discrete task variables^1,4–8^ (e.g., object categories^1^), the neural representations supporting OOD generalization in continuous task variables (e.g., driving speed) remain poorly understood.

It has been hypothesized that OOD generalization in continuous variables relies on low-dimensional, low-curvature neural population representations^2,9–11^. Studies in natural video perception^12^, primary visual cortex^13^, and full brain EEG recordings^14^ suggest possible low-dimensional, low-curvature manifold structures. However, direct evidence linking these geometric features to OOD generalization remains lacking. The underlying neural geometry supporting OOD generalization in continuous task variables remains unclear.

To address this question, we analyzed population responses in the mouse primary visual cortex (V1) to static grating stimuli of varying orientations (data from Stringer et al.^15^). We trained a decoder to extract orientation values from neural responses within a restricted orientation range and tested its generalization to novel, held-out orientation ranges. The results showed that V1 representations robustly support the decoder’s OOD generalization. Generalization errors correlated significantly with the dimensionality and curvature of the underlying neural manifold, suggesting that these geometric features are key indicators of a neural population representation’s OOD generalizability.

What learning objectives might give rise to this representational geometry? Supervised and reinforcement learning have both been shown to produce OOD-generalizable representations^16^, but they typically demand vast amounts of labeled data or extensive trial-and-error, which is often implausible in naturalistic settings. Input image reconstruction (e.g. variational autoencoders) can yield disentangled, generalizable features^1,17^; however, real-world visual inputs are continuous, dynamic streams rather than isolated, static images. Moreover, the geometric characteristics of the resulting manifolds (for example, their curvature) from these models have yet to be characterized. Thus, the plausible learning objectives that give rise to OOD-generalizable geometry remain elusive. Uncovering these objectives could not only shed light on how evolution has shaped the brain’s representational machinery but also offer high level intuition for designing training objectives in deep neural networks.

Inspired from input image reconstruction objective^1,17^, here we hypothesize that predictive learning— specifically, the ability to forecast future sensory inputs^18–21^—innately produces representational geometries that support OOD generalization. To test this, we studied PredNet, a deep network trained to predict the next frame in natural video sequences^19^. After training, mirroring the visual stimuli in Stringer et al. 2021^15^, we presented static gratings of varying orientations to the trained PredNet models, and characterized the geometry of its neural representations. We found that, as depth increases, PredNet’s layers form progressively lower-dimensional, lower-curvature manifolds and exhibit enhanced OOD generalization. These findings suggest that next-input prediction is a plausible learning objective for driving the hierarchical emergence of geometries conducive to robust OOD generalization in deep networks. Together, our results demonstrate a clear correlation between neural manifold dimensionality and curvature with OOD generalization, and highlight predictive learning as a plausible objective for acquiring such OOD-generalizable geometry.

## Results

### V1 representation supports out-of-distribution generalization

Out-of-distribution (OOD) generalization refers to the ability of a decoder to perform accurately on test data drawn from a novel domain. This capability depends not only on the decoder itself but also on the input data representation. Here, we investigate the extent to which neural population representations in the mouse primary visual cortex (V1) support OOD generalization.

We analyzed a previously published dataset collected by Stringer et al. (2021)^15^, in which mice passively viewed three types of grating stimuli with varying orientations (Figure 1A): rectangular gratings with fixed phase (six recording sessions), sine gratings with random phases (three sessions), and local gratings with fixed phase (three sessions). V1 population activity was recorded using two-photon calcium imaging. We refer to a dataset as a recording session’s neural population responses paired with the corresponding stimulus orientation.

**Figure 1.**
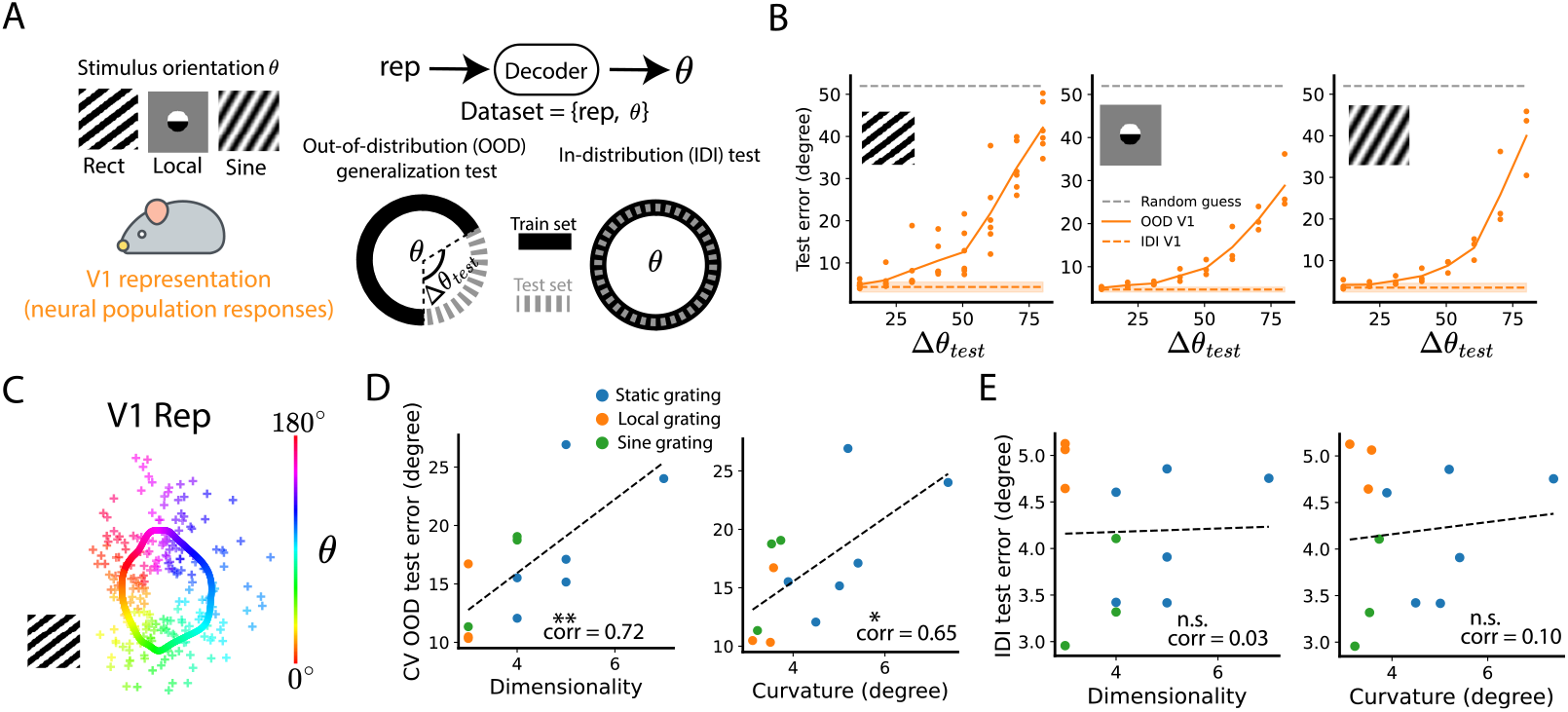
Decoders’ OOD generalization error correlates with the dimensionality and curvature of the underlying neural manifold. (**A**) We analyzed neural responses from mouse primary visual cortex (V1) to rectangular, sine, and localized grating stimuli of various orientations (data from Stringer et al. (2021)^15^). Each dataset includes V1 population responses and corresponding stimulus orientations. The dataset was split into train/test sets via two schemes: an OOD split, which partitions the dataset into two non-overlapping orientation ranges, and an IDI split, which samples data randomly. A decoder was trained to decode orientation from the neural responses. (**B**) OOD and IDI test errors. Each dot represents the result of a recording session at one Δ*θ*_*test*_. Lines connect session-averaged values. The orange dashed line indicates the mean IDI error; shaded bands denote the 95% confidence interval of the IDI test error mean (1.96 times standard error). Grey dashed line indicates the mean test error of a random guessing decoder (see Methods). (**C**) Neural manifolds were fit using Gaussian process regression (see Methods) and projected onto the PC1-PC2 plane for visualization. Crosses indicate test data excluded from fitting. (**D**) To have a single number representing the OOD generalization error of each recording session, we measured the cross-validation (CV) OOD test error (three folds with non-overlapping orientation ranges, see text and Methods). Manifold dimensionality was defined as the minimum number of principal components explaining 90% of the manifold variance. Curvature was estimated as the mean angle between adjacent tangent vectors on the manifold. Each dot is one recording session. Dashed lines indicate the best-fitting line (least mean squares). Pearson correlation coefficient (corr_ood) quantifies the relationship between OOD error and each geometric property. ***p<0.001, **p<0.01, *p<0.05, n.s.: not significant, two-sided Pearson correlation test^22^ (see Methods). (**E**) Same as panel (D), but y axis showing the IDI test error. Similar results but for CV OOD test error measured using k-nearest-neighbor (KNN) decoders are shown in SI Figure 2.

To evaluate the neural representation’s capability for OOD generalization, we split the dataset into training and test sets with non-overlapping label ranges (Figure 1A). Specifically, the test set comprised samples with orientation labels ranging from *θ*_*i*_ to *θ*_*i*_ + Δ*θ*_*test*_, where *θ*_*ii*_ was randomly chosen. The remaining data were assigned to the training set. We trained a decoder (circular ridge linear decoder, see Methods) to decode orientation values from the neural representations, and evaluated its performance on the test set. The test error (root mean squared error) served as a measure of OOD generalizability. We found that the generalization error gradually increased with increasing test range Δ*θ*_*test*_, but within the range tested, was consistently lower than that of a random decoder whose outputs are drawn from a uniform distribution (Figure 1B).

### OOD test error correlates with V1 neural manifold dimensionality and curvature

What properties of the neural representations predict OOD generalizability? The neural population representations are embedded in a high-dimensional space and are highly variable across trials. Despite this, recent advances in neural population geometry have hypothesized that neural population representations tend to be confined to a low-dimensional, nonlinear manifold^10^.

Based on this neural representation–geometry perspective, previous work has proposed that low dimensionality and low curvature of the manifold facilitate better OOD generalization^2,9–11^. To test this hypothesis, we inferred smooth neural manifolds from the data. Inferring manifolds from the original high-dimensional space (∼20 000 neurons) is challenging; therefore, we first projected the neural responses onto their first 200 principal components (PCs) using principal component analysis (PCA), capturing a large fraction of the variance explained (∼60%; see SI Figure 1). This yielded a dataset containing a 200-dimensional representation with corresponding grating-orientation labels. We then used Gaussian process regression to infer smooth manifolds (fitting r-squared > 0.9 for all recording sessions; see Methods). Specifically, we randomly split the dataset into training and test sets. The training set was used to fit a Gaussian process regression to establish a smooth mapping from orientation labels (0° to 180°) to the representational space, with “smoothness” constrained via a kernel function. We refer to this smooth mapping as the manifold. For visualization, we further projected the fitted manifold onto their first-two-PCs plane (PC1–PC2 plane), as shown in Figure 1C. The resulting plot demonstrates good alignment between the inferred manifold and the test data.

We discretized the inferred manifold into 300 equally spaced mesh points (see Methods). We defined manifold dimensionality as the minimum number of PCs required to explain the variance of these mesh points above a specified variance-explained-ratio threshold (var. ratio threshold); Manifold curvature as the mean angle between adjacent tangent vectors approximated by finite-difference vectors between neighboring mesh points (see Methods). Thus, for each recording session, we obtained one dimensionality value (at a fixed var. ratio threshold) and one curvature value (Figure 1D).

Do these dimensionality and curvature values correlate with the OOD generalization error? To answer this, we need a single scalar of OOD generalization for each recording session (rather than the different Δ*θ*_*test*_ in Figure 1B). Thus, we computed cross-validated (CV) OOD test error via three-fold cross-validation: a dataset was partitioned into three non-overlapping 60° orientation ranges; for each fold, a decoder was trained on the other two folds and tested on the held-out fold (ridge linear decoder: Figure 1D; k-nearest-neighbor decoder: SI Figure 2). The mean test error across all held-out folds defined the CV OOD test error, yielding one metric per recording session. As a baseline comparison, we also computed in-distribution (IDI) test errors by randomly splitting each dataset into training and test sets, and evaluating decoder performance on the test set after training on the training set.

**Figure 2.**
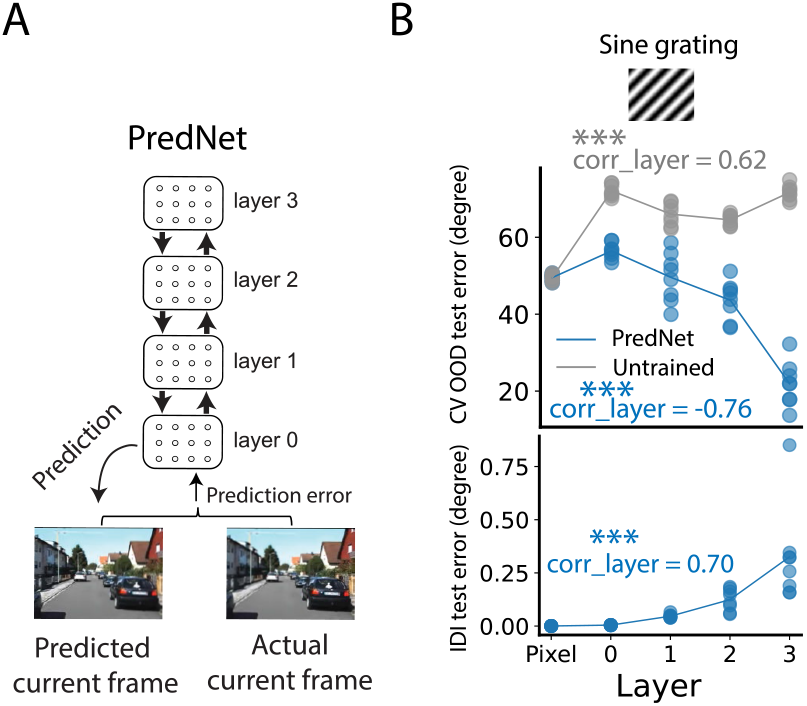
A deep neural network (PredNet) trained for natural video prediction forms an OOD-generalizable manifold in deeper layers. (**A**) PredNet comprises four hierarchical layers, each containing a representation module (R) for prediction and an error module (E) for computing prediction errors^19^. We trained eight PredNet models identically but with different random seeds on next-frame prediction of natural videos. As a baseline, we also inspected eight untrained PredNet models. (**B**) We presented sine grating stimuli with varying orientations to the PredNets, collected unit responses from the R modules of each layer (see similar results for E modules in SI Figure 4), and evaluated CV OOD test error (top) and IDI test error (bottom). Blue/grey dots represent the results of individual trained/untrained PredNet models. “Corr-layer” denotes the Pearson correlation between layer identity (with the pixel input denoted as -1) and test error. Statistical significance: ***p < 0.001, two-tailed Pearson correlation test^22^ (see Methods). Solid lines connect means.

We found that both manifold dimensionality and curvature were significantly correlated with CV OOD test error, but not with IDI test error (Figures 1D and 1E). Moreover, the significant correlations remained robust across different hyperparameter settings—such as the var. ratio threshold and the number of PCs (SI Figure 3). Together, these results support the hypothesis that manifold geometries (dimensionality and curvature) are indicative of OOD generalization.

**Figure 3.**
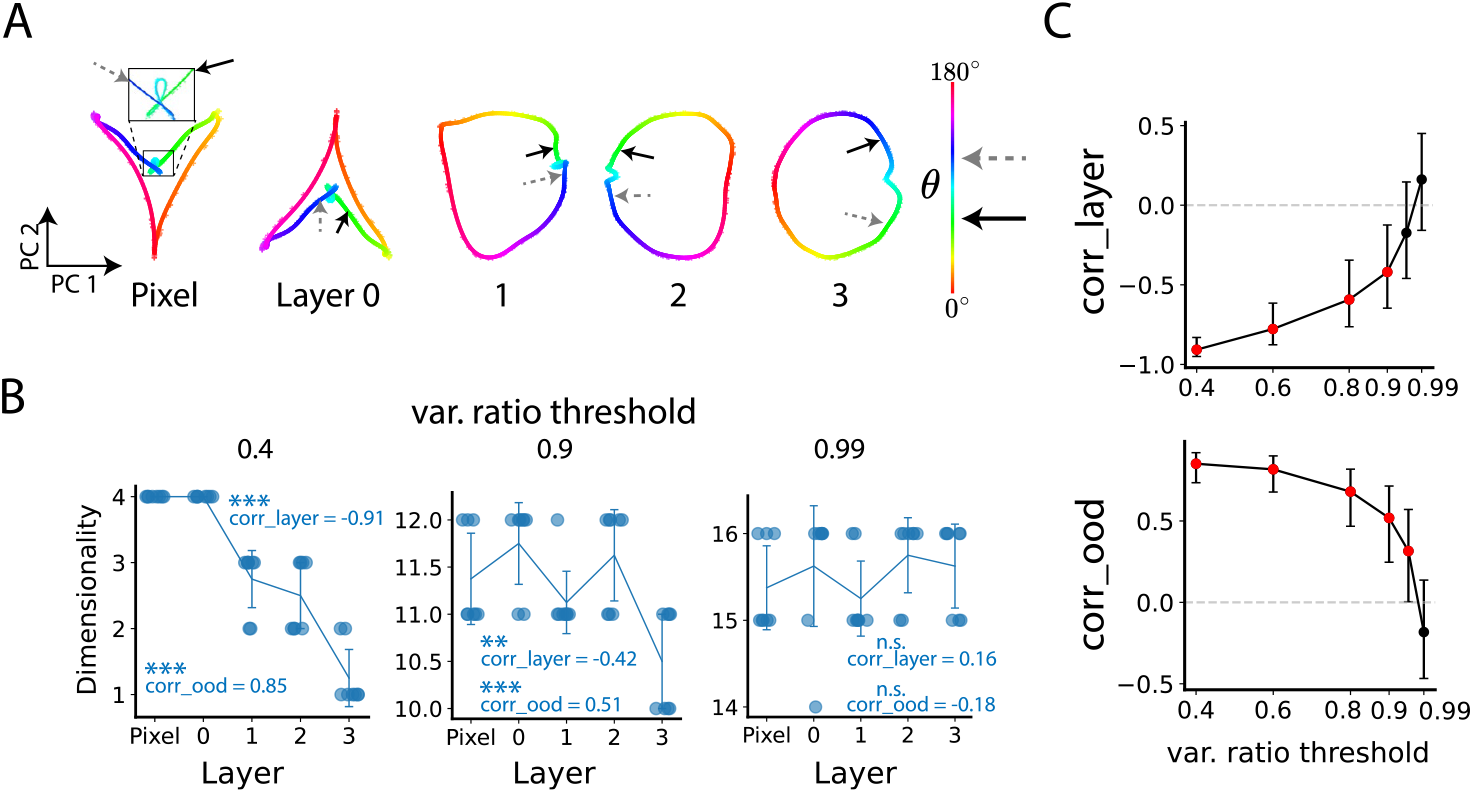
PredNet forms low-dimensional manifold. (**A**) We presented grating stimuli spanning various orientations and recorded R-module unit responses (see similar results for E modules in SI Figure 4). We then fit a smooth manifold to these responses using Gaussian process regression (see Methods). For visualization, we projected the manifold from one example trained PredNet onto its first two principal components. Crosses mark held-out test data not used during fitting; these points largely lie on the fitted manifold, likely reflecting PredNet’s noise-free responses and yielding high-quality fits (r-squared > 0.99). Two arrows indicate the representations of two fixed orientation values. (**B**) Manifold dimensionality is defined as the number of PCs (ranked by variance explained) required to exceed a specified variance ratio threshold. Panels show manifold dimensionality across different thresholds. Each dot represents a trained PredNet model. To improve visualization and reduce overlap, a small jitter was added along the x-axis (i.e. layer identity). Lines and error bars indicate the means and standard deviations across PredNet models. Corr_layer: Pearson correlation between dimensionality and layer identity (with the pixel layer labeled as -1). Corr_ood: Pearson correlation between dimensionality and CV OOD test error (see Figure 2B). Significance: ***p < 0.001; **p < 0.01; p < 0.05; n.s., not significant, two-sided Pearson correlation test (see Methods). (**C**) Correlations (Corr_layer and Corr_ood) of different var. ratio thresholds. Red dots indicate statistically significant correlations (p < 0.05). Dots and error bars representing the means and 95% confidence intervals (via Fisher z-transformation, see Methods).

### OOD-generalizable representations emerge in neural networks trained for next-frame natural video prediction

What learning objective enables a neural network to acquire OOD-generalizable representations? Predictive learning has been shown to give rise to multiple biologically plausible features^18,20,21^. Building on this, we hypothesize that predictive learning also promotes OOD-generalization. To test this, we examined PredNet—a deep neural network model inspired by predictive coding and trained to predict future frames in natural videos^19^.

PredNet comprises four hierarchical layers, each with a representation module (R) that generates predictions and an error module (E) that encodes prediction errors (Figure 2A). At each time step, activity propagates from the deepest layer to the first layer to produce a predicted frame. This prediction is then compared to the actual video input, and the resulting error is fed forward from the first layer back to the deepest layer to update internal representations for the next step. In our main analyses, we focused on the R modules, which encode predictive representations; results from the E modules, which convey prediction errors, are qualitatively similar and shown in SI Figure 4. Eight PredNet models were trained on videos recorded by a car-mounted camera (KITTI^23^) using different random seeds. As a baseline, we also evaluated eight untrained PredNet models.

**Figure 4.**
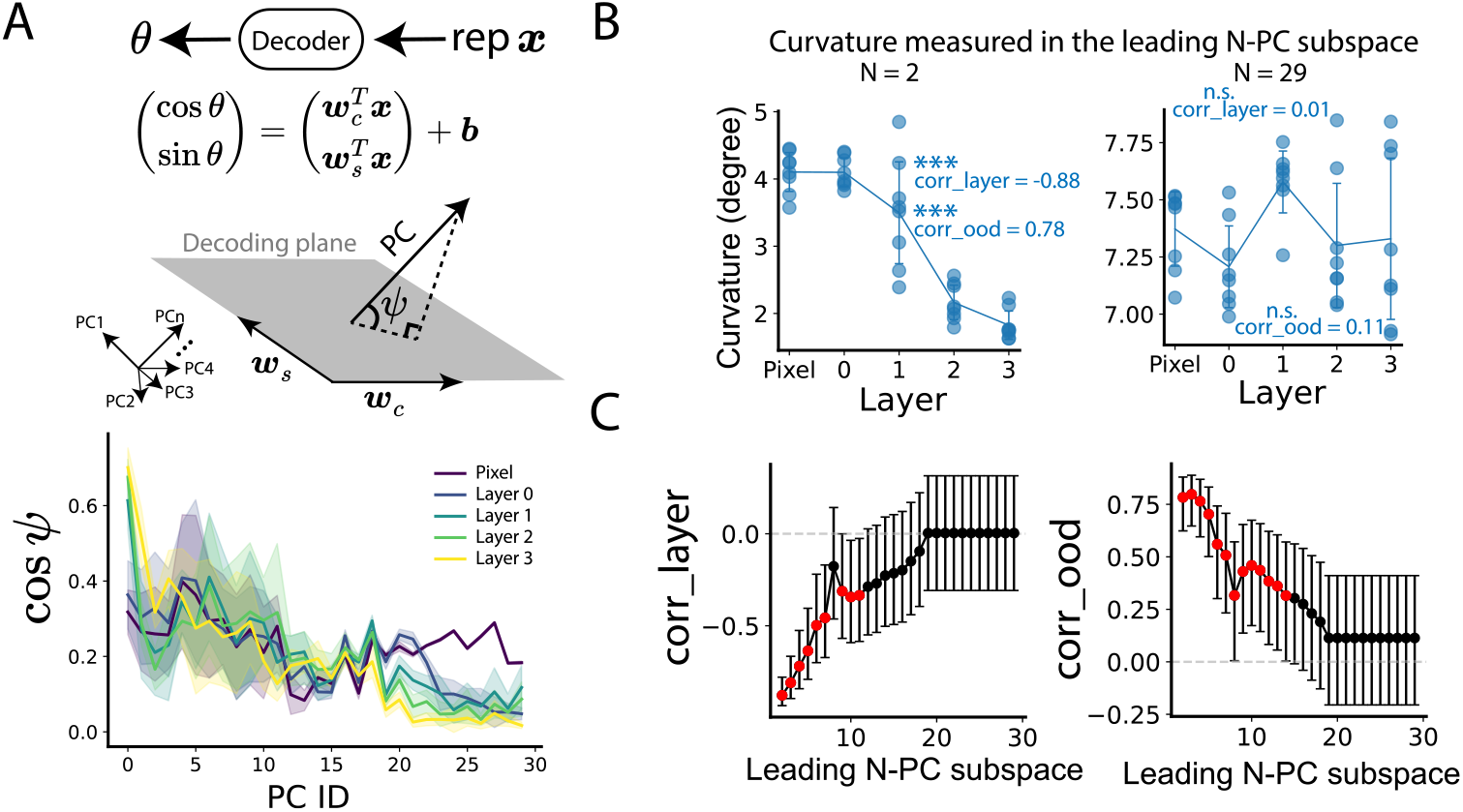
PredNet forms low-curvature manifolds. **(A)** A linear decoder extracts grating orientation *θ* from the representation ***x***. Geometrically, this decoder projects test data onto a “decoding plane” spanned by the two weight vectors ***w***_***s***_ and ***w***_***c***_, so only components within that plane contribute to the readout. To quantify each PC’s relevance. We computed the angle *ψ* between its axis and the decoding plane—lager values of cos *ψ* indicate greater relevance. Bottom: Lines and shaded bands show the mean ± standard deviation across eight independently trained PredNet models. (**B**) We projected the fitted manifold into subspaces spanned by the first N principal components and measured its curvature within each leading N-PC subspace. Illustrations are the same as Figure 3B. (**C**) Correlations between manifold curvature and layer identity (left) or CV OOD test error (right). Illustrations are the same as Figure 3C. All panels display unit representations from the R modules; similar results for E-module units are shown in SI Figure 4.

We presented sine grating stimuli with varying orientations to networks and collected unit responses across layers. Each dataset consisted of unit responses labeled by grating orientation. To reduce computational complexity, we reduced the dimensionality from about 50000 units in each module to 50 PCs, which captures a large fraction of data variance explained (≈ 1, SI Figure 1). Following the protocol in Figure 1D, we measured CV OOD test errors. We observed that CV OOD test error decreased in deeper layers of PredNet (R module in Figure 2B; E module in SI Figure 4A), indicating that deeper unit representations in PredNet are more OOD-generalizable. In contrast, the untrained PredNet models did not exhibit improvements in OOD generalization (R module in Figure 2B; E module in SI Figure 4A).

One might alternatively attribute the hierarchical reduction in CV OOD test error to improved in-distribution decoding rather than genuine OOD generalization. To assess this, we measured IDI test errors using random train–test splits (as in Figure 1D, right panel). Contrary to that interpretation, IDI test error slightly increased in deeper layers (PredNet R modules in Figure 2B, bottom panel; E modules in SI Figure 4A), indicating that in-distribution decoding performance worsens with depth. This decline likely reflects the certain randomness of PredNet’s learned weight values, which “disrupt” the otherwise noiseless pixel representations. Nonetheless, IDI test errors remained consistently low across all layers (< 1°), indicating minimal decoding difficulty in the in-distribution case. Together, these results demonstrate that the reduction in CV OOD test error cannot be explained by IDI decoding performance, supporting the conclusion that predictive learning in PredNet yields genuinely OOD-generalizable representations.

### PredNet forms low-dimensional manifolds in deep layers

What representation geometry underlies PredNet’s OOD-generalizable representations? To uncover this, we inferred neural manifolds from PredNet’s unit responses to sine-grating stimuli. For each layer, we collected unit activations and projected them onto the first 50 PCs for computational efficiency and numerical stability (capturing ∼100 % of the variance, see SI Figure 1). We then fit a smooth manifold to these projections using Gaussian process regression (r-squared > 0.99 for all modules). Example manifolds, visualized in the PC1–PC2 plane, are shown in Figure 3A (R modules) and SI Figure 4B (E modules). Visual inspection suggests that these manifolds become progressively lower dimensional and less curved in deeper layers.

To quantify this trend, same as neural data (Figure 1D, left panel), we discretized each manifold into 300 evenly spaced mesh points and defined its dimensionality as the minimum number of PCs required to exceed a given var. ratio threshold. Across most thresholds below 0.9, deeper PredNet layers exhibited significantly lower dimensionality than early layers (Figures 3B and 3C upper panels for R modules; SI Figure 4C for E modules). Moreover, dimensionality correlated strongly with CV OOD test error (Figure 3C, lower panel; SI Figure 4C), suggesting that PredNet formed lower-dimensional representations that are indicative of OOD generalization.

### OOD-generalizable information in PredNet predominantly lie within the low-dimensional subspace defined by the leading principal components

Unlike the robust correlation observed in V1 data (SI Figure 3), however, the relationship between manifold dimensionality and OOD generalization in PredNet breaks down as the var. ratio threshold approaches 1 (e.g., 0.99; Figure 3B, right panel for R modules; SI Figure 4C for E modules). At these high thresholds, dimensionality no longer correlates with layer depth (Figure 3C, top panel for R modules; SI Figure 4C for E modules) or with CV OOD error (Figure 3C, bottom panel for R modules; SI Figure 4C for E modules). This result suggests a conjecture that including nearly all variance incorporates high-order “noise” PCs that do not contribute to OOD performance; instead, the subspace spanned by the leading PCs is what truly matters for generalization.

To test this conjecture, we quantified each PC’s relevance to orientation decoding. A linear decoder predicts grating orientation by projecting test data onto a decoding plane defined by its two weight vectors (Figure 4A); only the component within this plane contributes to decoding. We therefore measured cos *ψ* where *ψ* is the angle between each PC axis and the decoding plane (see Methods). Consistent with our conjecture, the leading PCs closely align with the decoding plane across all PredNet layers (Figure 4A and SI Figure 4D), indicating that OOD-generalizable information primarily resides in PredNet’s low-dimensional subspace formed by leading PCs.

### PredNet forms low-curvature manifold in leading N-PC subspace

In addition to dimensionality, we examined curvature as another key geometric property. We meshed the manifold into 300 points and defined curvature as the angle between adjacent tangent vectors. In the high-dimensional space (e.g., 29 dimensions), curvature did not correlate with PredNet layer depth or CV OOD generalization error (Figure 4B, right panel and SI Figure 4E). However, as noted above, high-dimensional representations may include “noise” dimensions. To address this, we projected the 300 mesh points into subspaces spanned by the first N principal components (leading N-PC subspaces). For small N, deeper PredNet layers exhibited lower-curvature manifolds, and curvature within these leading N-PC subspaces correlated with CV OOD generalization error (Figure 4C and SI Figure 4E). Together, these results indicate that low curvature within leading N-PC subspaces is a geometric signal of OOD-generalizable representations in PredNet. These curvature findings (Figure 4), together with the manifold dimensionality results (Figure 3), indicate that training a neural network solely on stimulus prediction yields geometrically OOD-generalizable representations.

## Discussion

OOD generalization is a hallmark of intelligent systems, yet the specific geometric features of neural population representations that support it—especially on continuous variable domains (e.g., orientation or driving speed)—remain poorly understood. Here, we demonstrate that both manifold dimensionality and curvature correlate with a representation’s ability to generalize out-of-distribution (Figure 1).

Moreover, a deep network trained to predict the next frame in natural video (PredNet^19^) develops increasingly OOD-generalizable representations in its deeper layers (Figures 2, 3, and 4), suggesting that predictive learning alone—without explicit supervisory labels—can give rise to representational geometry conducive to robust generalization.

Although both PredNet and mouse V1 qualitatively exhibit low dimensionality and low curvature (Figures 1D, 3 and 4), closer examination reveals systematic differences. PredNet’s low-dimensional and low-curvature representation geometry is evident only when considering the submanifold that accounts for a limited portion of the total variance (Figures 3C and 4C), whereas mouse V1 maintains these geometric properties across a wide range of var. ratio thresholds (SI Figure 3). This implies a possibly more robust representation in the biological neural system^24,25^. Several factors may explain this difference between V1 and PredNet. First, biological neural activity is inherently noisy, and training neural networks with noise has been shown to promote representation robustness^26–28^ and straighten the neural manifold^29^. Thus, trial-to-trial variability in V1 may enhance manifold generalizability. Second, the visual cortex must support many functions—recognition, motion detection, and more—whereas PredNet is optimized solely for prediction; multitask learning is known to improve representational generality^16,30^. Third, input statistics differ: PredNet views static gratings, but the natural visual stream in mice is constantly modulated by eye and head movements, introducing richer dynamics that may shape manifold geometry. Lastly, the architectural distinctions between brains and PredNet may also contribute to their representational differences^20,31^. Unraveling the underlying factors that lead to differences in representations between the brain and deep neural networks—and even developing novel methods to narrow these gaps—are important directions for future work.

Although our analysis here is restricted to simple static gratings, the framework may extend naturally to more complex, multidimensional stimuli^32,33^. For instance, the dSprites “Disentanglement” dataset comprises objects parameterized by continuous size, position, and orientation variables etc^33^. Neural responses to these stimuli may form intrinsically high-dimensional manifolds (for example, a surface^34^) rather than one-dimensional curves (Figure 1C). Nevertheless, one can still measure dimensionality and curvature in these higher-dimensional manifolds to test whether these geometric features signify OOD performance. Beyond synthetic images, the framework of this paper may also extend to natural images and videos. Indeed, recent work shows that primary visual cortex encodes temporal sequences in relatively straight, low-curvature trajectories, hinting at a link to temporal generalization^13^. Understanding the OOD-generalization geometry for more complex and natural stimuli remains a future direction.

PredNet exhibits a hierarchical improvement in both OOD generalization (Figure 2) and representational geometry (Figures 3 and 4). Does the brain likewise refine its OOD-generalizable geometry across the visual hierarchy^9^? Studies of discrete-variable generalization—e.g., object recognition across novel contexts—suggest that higher visual areas in macaques may support more generalizable representations^1,7^. In contrast, investigations of continuous-variable generalization are relatively scarce. Does the OOD generalization of continuous variables (e.g., orientation) also improve in higher-order visual hierarchies? Single-unit recordings support this by demonstrating that neurons in downstream areas show more disentangled tuning to object features^2,35,36^. However, direct measures of the correlation between neural population activity across different hierarchies and its OOD generalization are still lacking. Characterizing how manifold geometry and generalization evolve through cortical hierarchies remains an interesting avenue for future exploration. Together, our results establish a clear correlation between manifold geometry and OOD generalization (Figure 1), opening new possibilities for understanding generalizable neural representations.

## Methods

### V1 neural data and the stimuli used

Experimental data were obtained from Stringer et al. (2021)^15^. Neural activity was recorded via multi-plane two-photon calcium imaging in primary visual cortex (V1) of awake, head-fixed mice running freely on an air-floating ball, capturing approximately 20,000 neurons per session. Fluorescence signals were deconvolved to infer spike timing, and stimulus-evoked responses were averaged over two consecutive 650 ms time bins following stimulus onset for downstream analyses.

We analyzed responses to three types of grating stimuli (Figure 1A): Rect: static rectangular gratings with a fixed phase (six recording sessions); Local: localized static gratings with a fixed phase (three recording sessions); Sine: static sine gratings with randomized phases (three recording sessions). Each session comprised 3,500–5,500 trials, where each trial consisted of a single grating presentation with directions drawn at random. Because a grating at direction *ϕ* is perceptually similar to *ϕ* + 180^°^, we defined the orientation *θ* as *ϕ* modulo 180^°^ and only considered orientations as labels in this paper.

### Circular ridge regression and circular KNN regression

Orientation labels *θ*, defined on the circular interval [0^°^, 180^°^), were first embedded on the unit circle as (cos*θ*, sin*θ*). For circular ridge regression, we then fit two ridge models—one for the sine component and one for the cosine component—using the same L2-penalty coefficient α. We select α by two-fold cross-validation over the grid {0.1, 1, 10, 100}. At prediction, the model returns *ŷ*_*sin*_ and *ŷ*_*cos*_, which we convert back to an angular estimate via an arctangent transformation. This procedure, referred to as circular ridge regression, avoids discontinuities at the 0^°^*/*180^°^ boundary by operating in the continuous sine–cosine domain.

Besides, we also implemented a circular k-nearest neighbor (KNN) regressor (SI Figure 2) by training two independent KNeighborsRegressor models (from scikit-learn^37^) on the sine and cosine embeddings, respectively. For each test sample, each regressor returns the mean of its k = 20 nearest neighbors in the training set for the corresponding component; these two averaged values are then recombined into an orientation estimate via the same arctangent-based conversion.

### Out-of-distribution (OOD) test error and in-distribution (IDI) test error

A dataset consists of neural responses paired with grating-orientation labels.

In Figure 1B, we considered the OOD test error for different choices of Δ*θ*_*test*_. Specifically, data with labels in the range (*θ*_*i*_, *θ*_*i*_ + Δ*θ*_*test*_) were assigned to the test set, and the rest to the training set. We trained a decoder (circular ridge) on the training set and compute the root-mean-squared error—using circular difference—on the test set, yielding one OOD test error per (*θ*_*i*_, Δ*θ*_*test*_) pair. For each fixed Δ*θ*_*test*_, we sampled three *θ*_*i*_ values at random and average their OOD test errors; this average is plotted as a single dot in Figure 1B.

We also evaluated the IDI test error in Figure 1B. The computation is identical to that for the OOD test error, except that the training and test sets are randomly split (train: test = 0.66:0.33). The dashed line shows the mean IDI error across all recording sessions and Δ*θ*_*test*_ (although it should not depend on Δ*θ*_*test*_); and the shaded band represents the 95% confidence interval (± 1.96 × standard error).

In Figures 1D, 2B, 3C, and 4C, we reported the cross-validated (CV) OOD test error as a single measure of a dataset’s OOD generalizability. A dataset was split into three non-overlapping folds, each covering 60^°^ of orientation. A decoder was trained on two folds and tested on the held-out fold, yielding three OOD errors that were then averaged to give the CV OOD test error.

For comparison to CV OOD test error in Figures 1E and 2B, we computed the IDI test error using a similar procedure, except that the splits are random rather than by label range. Thus, the IDI test error of a dataset is the average of three IDI errors from three random splits.

### Test error of a random guess decoder

Consider the extreme case in which the representation is so poor that the decoder can only guess at random. We refer to this as a random-guess decoder (Figure 1B), whose output draws from a uniform distribution over [0^°^, 180^°^). Consequently, the circular difference between prediction and ground truth, 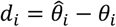, follows a uniform distribution on −90^°^ to 90^°^. The test error is the root-mean-squared-error 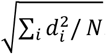. It then follows that 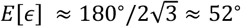 (using a delta-method approximation^38^ for the square root). This average error is indicated by the grey dashed line in Figure 1B.

### Gaussian process regression (GPR) for manifold fitting

A representation (pixel values, V1 neural responses or PredNet unit activations) is a high-dimensional vector (e.g. ∼ 20000 neurons in V1), which can be both computationally expensive and numerically unstable to process. To address this, we first applied principal component analysis (PCA) to reduce each representation to *M* dimensions (V1: *M* = 200, cumulative variance explained ratio ≈ 0.6; PredNet: *M* = 50, cumulative variance explained ratio ≈ 1; see SI Figure 1), denoted as ***x***. We then split the dataset 4:1 into training and test sets and fitted a Gaussian process regression (GPR) on the training data, using the orientation label *θ* as independent variable and the M-dimensional vector *x* as the dependent variable^34,39–42^.

Specifically, prior to fitting, *x* was standardized to zero mean and unit variance, yielding 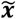, whose expected value 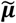 was modeled as an N-independent Gaussian process (*N = M*) with the periodic kernel function to control the “smoothness” of 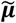 over label *θ*

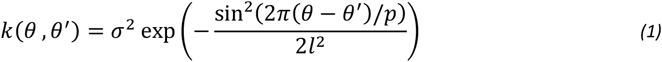

with a period *p* = *π*. Hyperparameters were optimized by maximizing the log-likelihood of the joint Gaussian distribution 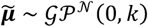. Finally, 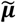 was unstandardized back to ***µ***.

In vanilla Gaussian process regression, constructing an *K* × *K* kernel matrix where *K* is the number of data points. This could be quite computationally expansive, for example, considering about 4000 data points in each of the V1 recording sessions. Therefore, we adopted the inducing variance method^41^. It approximates the training data labels by a smaller set of inducing points. In this paper, inducing points were initialized as a randomly sampled subset of the original training labels (30 inducing points), and were optimized during the optimization of Gaussian process regression. The Gaussian process regression with inducing variables method is implemented using the Python GPflow package^40^. For more details and computer codes see Ye et al. (2024)^34^.

After fitting the manifold, we applied a one-nearest-neighbor decoder to assess its fitting quality. Specifically, for each held-out test point, we identified the closest mesh point on the manifold and assigned that mesh point’s label as the decoded value. Repeating this across the entire test set yielded an r-squared score that quantifies the goodness of the GPR fit. The resulting r-squared score for all V1 recording sessions and PredNet modules are larger than 0.9.

To visualize the fitted manifold (Figures 1C and 3A), we first sampled 300 equal-spacing label values and used GPR to compute their corresponding points on the manifold. We then performed PCA on these 300 points and projected them onto the plane defined by the first two PCs for display. Held-out test data—not used during fitting—were projected into the same PC plane (crosses in Figures 1C and 3A), allowing a direct visual comparison between the inferred manifold and the raw data.

### Manifold dimensionality and curvature

We first meshed the manifold into 300 points and applied PCA to this set. By computing the cumulative variance explained by each PC, we define the manifold’s dimensionality as the smallest number of PCs required to exceed a specified variance-explained ratio threshold (var. ratio threshold).

The curvature is the angular difference of two adjacent tangent vectors. We used the same definition as the one used in Henaff et al. 2019^12^

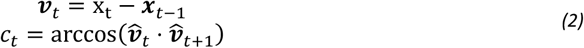

where *x*_*t*_ is the *t*-th mesh point on the manifold, 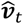 is a normalized ***v***_*t*_ (unit length). The manifold’s curvature is the mean of curvatures across 299 mesh points (excludes the boundary one).

### Pearson correlation and statistical test

Throughout this paper, the term “correlation” refers to the Pearson correlation coefficient. P-values were computed using SciPy’s scipy.stats.pearsonr function, which performs a two-tailed test under the null hypothesis that the true correlation *r* = 0. The null distribution was *f*(*r*) = (1 − *r*^2^)^*s/*2−2^*/ B*(1*/*2, *n/*2 − 1) where *n* is the number of sample points, and *B* is the beta function. The observed correlation coefficient was compared against this distribution to yield a p-value, indicating the probability of observing a correlation at least as extreme as the one obtained purely by chance under the assumption of no linear relationship between the variables.

To estimate the 95 % confidence interval around each correlation *r*, we applied the Fisher z-transformation from *r* to *z* which follows approximately a normal distribution. Specifically, the observed correlation *r* was converted to *z* = arctanh(*r*). Under the assumption of independent observations, z has a standard error of 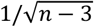, where *n* is the sample size. We then identified the critical value from the standard normal distribution corresponding to the desired 95 % confidence (i.e., 1.96) and computed the lower and upper bounds in z-space as 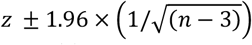. Finally, these bounds were back-transformed to the correlation scale via *r* = tanh(*z*) to yield the lower and upper confidence limits around the original *r*. These are shown in Figures 3C and 4C.

### PredNet architecture and training

We employed PredNet (code: https://github.com/coxlab/prednet), a four-layer predictive-coding-inspired network for next-frame natural video prediction^19^. Each layer comprises four modules: a representation module (*R*_*l*_), a prediction module (*Âl*), a target module (*A*_*l*_) and an error module (*E*_*l*_). Since R modules are directly used for making predictions, we focus on the analysis of R modules in the main figures, while presenting the results of the same analysis on the E module in SI Figure 4.

At each time step *t*, PredNet updates the *R*_*l*_ modules from the deepest to the first layer to generate a prediction *Â* _0_ of the next frame. This prediction is compared to the true frame *A*_0_ to compute the error *E*_0_, which is then fed forward to deeper layers to inform the subsequent prediction.

PredNet’s weights were optimized for next-frame prediction using videos recorded by the car-mounted cameras in the KITTI dataset^23^. Specifically, videos were center-cropped and downsampled to 128 × 160 pixels, segmented into 4,212 ten-frame sequences, and split into 4,114 training, 83 validation and 15 test sequences. The objective is to minimize the mean prediction error:

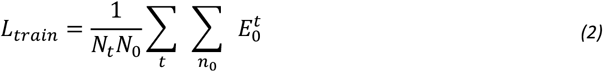

where *N*_*t*_ is the number of time steps, *N*_0_ is the number of *E*_0_ units. We optimized PredNet weights with the Adam optimizer. Eight PredNet models were trained with identical settings but different random seeds; as a control, we also analyzed eight randomly-initialized (untrained) PredNet models (Figure 2B). For more details see Lotter et al. (2017, 2020)^19,20^.

### PredNet responses to sine grating stimuli

We generated 800 sine-grating stimuli (128 × 160 pixels) with random orientations, a fixed phase, and a spatial period of 30 pixels. Each image was repeated four times along the temporal axis to create a four-frame sequence, which was then presented to PredNet. For each sequence, unit activations from time steps 2–4 were averaged to yield an orientation-specific representation.

Because each PredNet module contains roughly 50,000 units, we applied PCA to reduce these representations to their first 50 PCs (cumulative variance explained ratio ≈ 1, see SI Figure 1). This 50-dimensional vector was used for analyses—e.g., computing CV OOD test error in Figure 2B, or fitting manifold (see Methods: Gaussian Process Regression for Manifold Fitting).

### Angles between the decoding plane and PCs

A linear decoder estimates orientation by projecting a test data point onto a two-dimensional decoding plane defined by two weight vectors (Figure 4A; *see Methods: Circular ridge regression and circular KNN regression*). To quantify how much each PC contributes to decoding, we measure the angle—or more directly, the cosine of the angle—between each PC and this plane.

We first fit a circular ridge decoder using the same procedure as *Methods: Out-of-distribution (OOD) test error and in-distribution (IDI) test error*. Specifically, the data with labels in (*θ*_*i*_, *θ*_*i*_ + 60^°^) were held out as the test set; the rest formed the training set; where *θ*_*i*_ was drawn in random for each module.

After training, we extracted the two weight vectors, normalized them, and orthogonalized them via the Gram–Schmidt process to obtain an orthonormal basis (***u***_1_, ***u***_2_). Note the test data were not used.

We then performed PCA on the full dataset to obtain PC vectors ***P***. Each PC’s projection onto the decoding plane was computed as: ***P***_*proj*_ = (***P*** ⋅ ***u***_1_)***u***_1_ + (***P*** ⋅ ***u***_2_) ***u*** _2_. The cosine value between ***P*** and the plane is given by ***P***_*proj*_*/* ‖***P*** ‖ where ‖ ⋅ ‖ denotes the Euclidean norm.

## Author contributions

Conceptualization: ZY, RW Methodology: ZY, RW Investigation: ZY Supervision: RW

Writing: ZY, RW

## Data and materials availability

The analysis code is available at https://github.com/AgeYY/ood-geom-v1-net.git

## Supplementary Information

**SI Figure 1.**
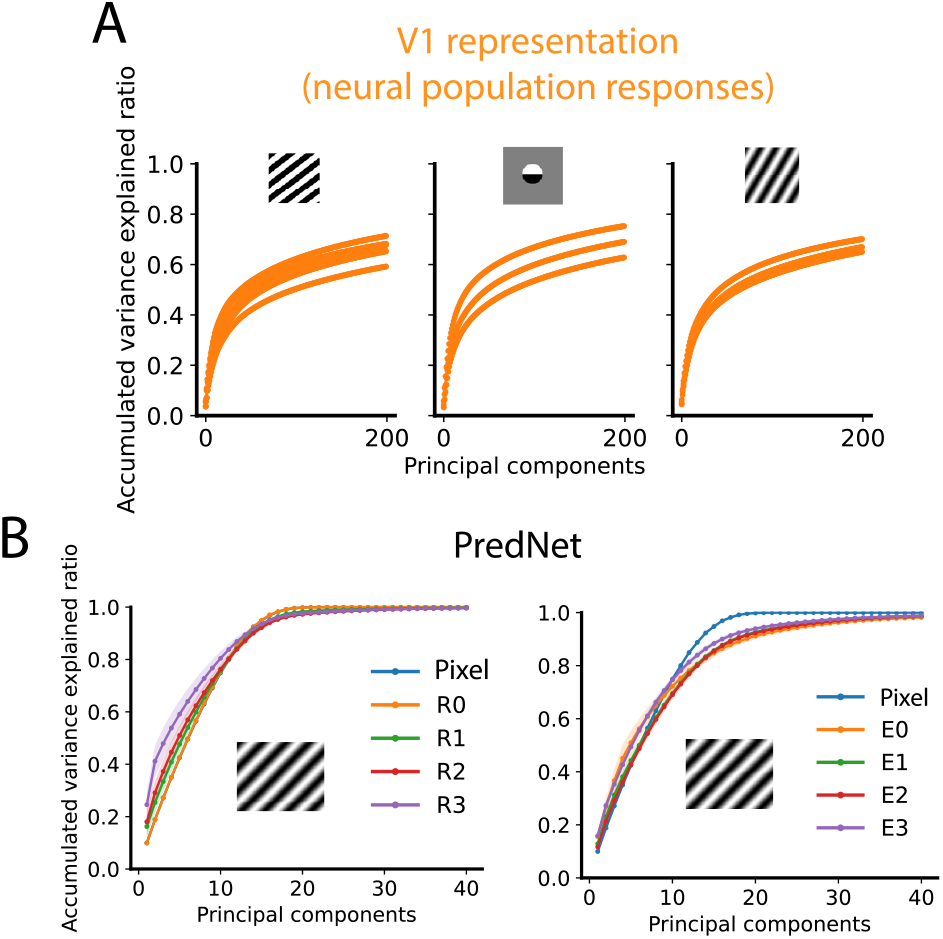
Cumulative variance explained ratio. (**A**) Each line represents the cumulative variance explained ratio of a single recording session. (**B**) Responses of PredNet units to 800 sine-grating stimuli with varied orientations (fixed phase) were analyzed by PCA (see Methods). Lines and shaded bands denote the mean ± one standard deviation across eight independently trained PredNet models.

**SI Figure 2.**
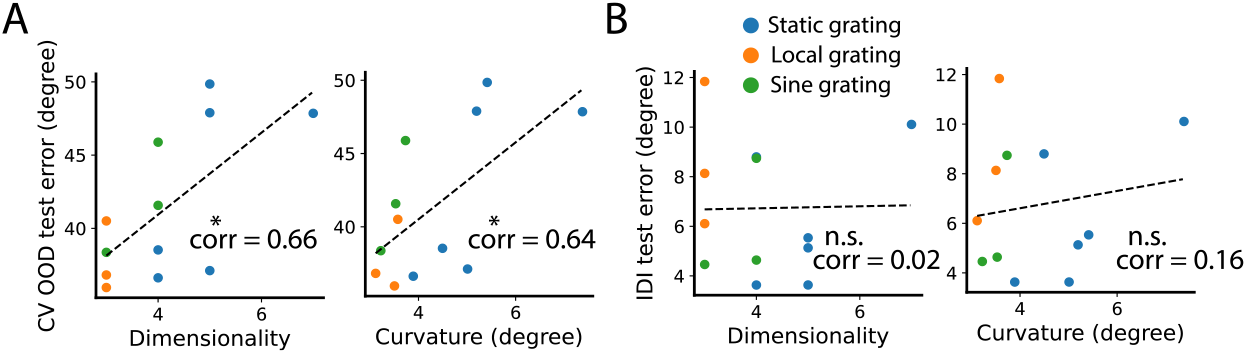
CV OOD test error—but not IDI test error—of V1 neural representations correlates with manifold dimensionality and curvature. (**A, B**) Plotting conventions follow Figures 1D and 1E, except that a circular k-nearest neighbors (KNN) decoder was used (see Methods).

**SI Figure 3.**
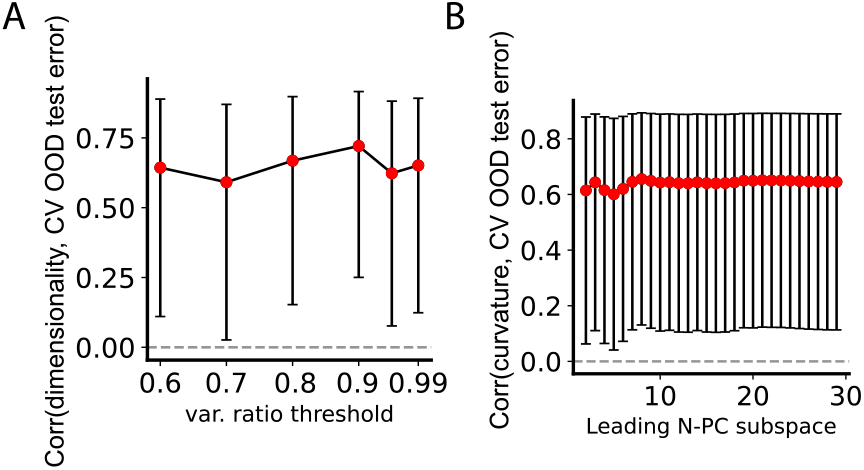
Correlations between V1 manifold geometry and CV OOD test error remain robust across different hyperparameter settings. (**A**) For each variance-explained ratio threshold (var. ratio threshold), we computed the Pearson correlation between manifold dimensionality and CV OOD test error across recording sessions. Each point represents the mean correlation; error bars denote the 95% confidence interval (see Methods). Red points indicate statistically significant correlations (p < 0.05, two-sided Pearson correlation test; see Methods). The result for the 0.9 threshold is shown in Figure 1D. (**B**) For each session, we fitted a neural manifold, discretized it into 300 mesh points, and projected those points onto the subspace defined by the first N principal components. At each choice of N, we calculated the Pearson correlation between manifold curvature and CV OOD test error across recording sessions. Figure 1D shows the result when N = 200. Point and error-bar conventions are the same as in panel (A).

**SI Figure 4.**
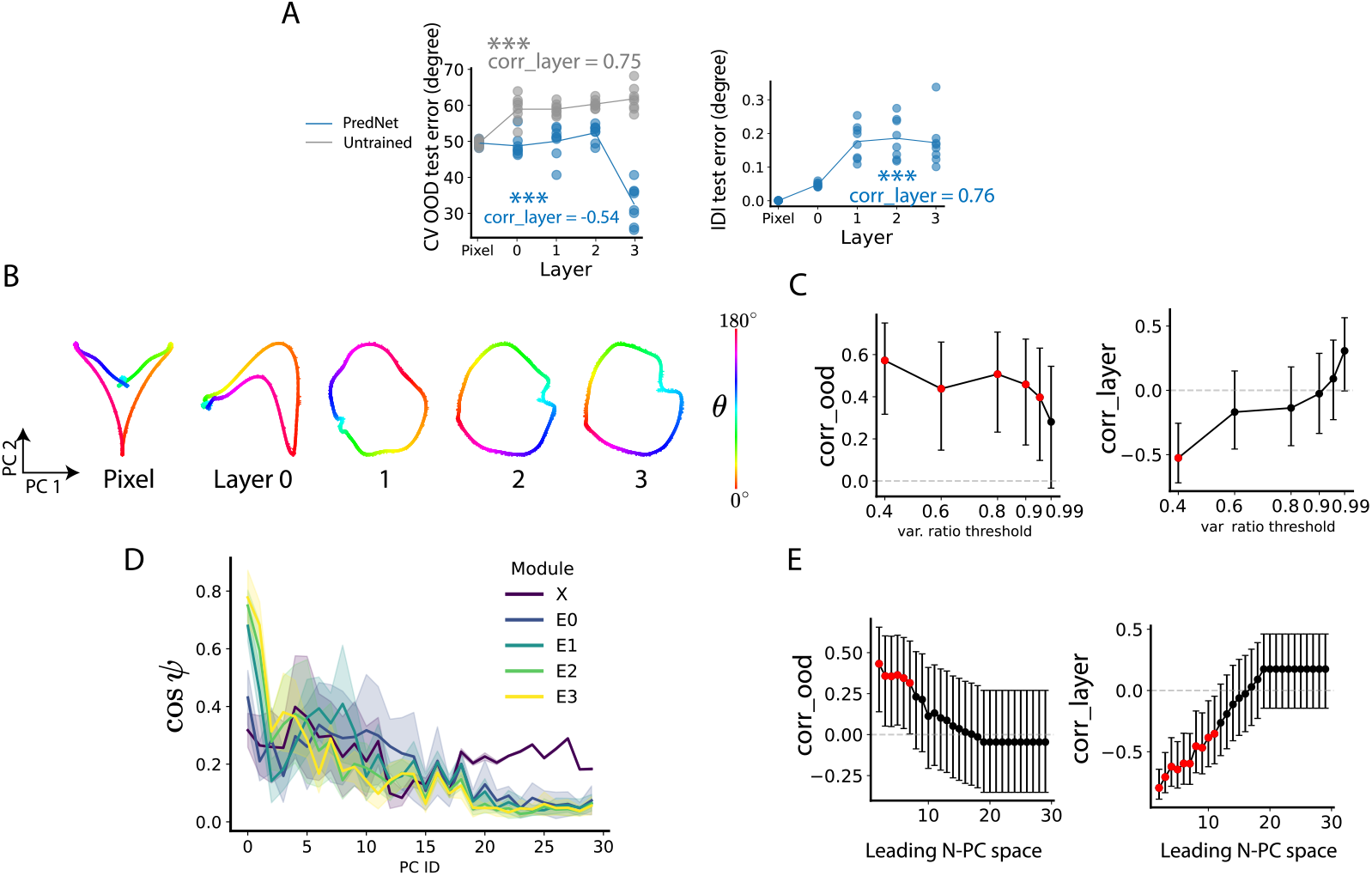
PredNet forms OOD-generalizable representations in deeper layers. (**A**) Same as Figures 2, 3, and 4, but for PredNet E modules.

## References

1. Li, Q., Sorscher, B. & Sompolinsky, H. Representations and generalization in artificial and brain neural networks. Proceedings of the National Academy of Sciences 121, e2311805121 (2024) DOI: 10.1073/pnas.2311805121.

2. Higgins, I., Racanière, S. & Rezende, D. Symmetry-Based Representations for Artificial and Biological General Intelligence. Frontiers in Computational Neuroscience 16, 1–16 (2022) DOI: 10.3389/fncom.2022.836498.

3. Ostojic, S. & Fusi, S. Computational role of structure in neural activity and connectivity. Trends in Cognitive Sciences 28, 677–690 (2024) DOI: 10.1016/j.tics.2024.03.003.

4. Kubilius, J., Kar, K., Schmidt, K. & DiCarlo, J. Can Deep Neural Networks Rival Human Ability to Generalize in Core Object Recognition? 2018 Conference on Cognitive Computational Neuroscience 3, (2018) DOI: 10.32470/ccn.2018.1234-0.

5. Bernardi, S. et al. The Geometry of Abstraction in the Hippocampus and Prefrontal Cortex. Cell 183, 954-967.e21 (2020) DOI: 10.1016/j.cell.2020.09.031.

6. Madan, S. et al. When and how convolutional neural networks generalize to out-of-distribution category–viewpoint combinations. Nature Machine Intelligence 4, 146–153 (2022) DOI: 10.1038/s42256-021-00437-5.

7. Sorscher, B., Ganguli, S. & Sompolinsky, H. Neural representational geometry underlies few-shot concept learning. Proceedings of the National Academy of Sciences of the United States of America 119, 1–12 (2022) DOI: 10.1073/pnas.2200800119.

8. Cohen, U., Chung, S. Y., Lee, D. D. & Sompolinsky, H. Separability and geometry of object manifolds in deep neural networks. Nature Communications 11, 1–13 (2020) DOI: 10.1038/s41467-020-14578-5.

9. DiCarlo, J. J., Zoccolan, D. & Rust, N. C. How Does the Brain Solve Visual Object Recognition? Neuron 73, 415–434 (2012) DOI: 10.1016/j.neuron.2012.01.010.

10. Chung, S. & Abbott, L. F. Neural population geometry: An approach for understanding biological and artificial neural networks. Current Opinion in Neurobiology 70, 137–144 (2021) DOI: 10.1016/j.conb.2021.10.010.

11. Bengio, Y., Courville, A. & Vincent, P. Representation Learning : A Review and New Perspectives. Preprint at 10.48550/arXiv.1206.5538 (2012) DOI: 10.48550/arXiv.1206.5538.

12. Hénaff, O. J., Goris, R. L. T. & Simoncelli, E. P. Perceptual straightening of natural videos. Nature Neuroscience 22, 984–991 (2019) DOI: 10.1038/s41593-019-0377-4.

13. Hénaff, O. J. et al. Primary visual cortex straightens natural video trajectories. Nature Communications 12, (2021) DOI: 10.1038/s41467-021-25939-z.

14. Sheahan, H., Luyckx, F., Nelli, S., Teupe, C. & Summerfield, C. Neural state space alignment for magnitude generalization in humans and recurrent networks. Neuron 109, 1214-1226.e8 (2021) DOI: 10.1016/j.neuron.2021.02.004.

15. Stringer, C., Michaelos, M., Tsyboulski, D., Lindo, S. E. & Pachitariu, M. High-precision coding in visual cortex. Cell 184, 2767-2778.e15 (2021) DOI: 10.1016/j.cell.2021.03.042.

16. Johnston, W. J. & Fusi, S. Abstract representations emerge naturally in neural networks trained to perform multiple tasks. Nat Commun 14, 1–18 (2023) DOI: 10.1038/s41467-023-36583-0.

17. Higgins, I. et al. beta-VAE: Learning Basic Visual Concepts with a Constrained Variational Framework. in Proceedings of the 5th International Conference on Learning Representations (ICLR) (2017). URL: https://openreview.net/forum?id=Sy2fzU9gl.

18. Rao, R. P. N. & Ballard, D. H. Predictive coding in the visual cortex: a functional interpretation of some extra-classical receptive-field effects. Nat Neurosci 2, 79–87 (1999) DOI: 10.1038/4580.

19. Lotter, W., Kreiman, G. & Cox, D. Deep Predictive Coding Networks for Video Prediction and Unsupervised Learning. Preprint at 10.48550/arXiv.1605.08104 (2017) DOI: 10.48550/arXiv.1605.08104.

20. Lotter, W., Kreiman, G. & Cox, D. A neural network trained for prediction mimics diverse features of biological neurons and perception. Nat Mach Intell 2, 210–219 (2020) DOI: 10.1038/s42256-020-0170-9.

21. Ye, Z., Wessel, R. & Franken, T. P. Brain-like border ownership signals support prediction of natural videos. iScience 28, (2025) DOI: 10.1016/j.isci.2025.112199.

22. Student. Probable error of a correlation coefficient. Biometrika 6, 302–310 (1908) DOI: 10.1093/biomet/6.2-3.302.

23. Geiger, A., Lenz, P., Stiller, C. & Urtasun, R. Vision meets robotics: The KITTI dataset. Int. J. Rob. Res. 32, 1231–1237 (2013) DOI: 10.1177/0278364913491297.

24. Safarani, S. et al. Towards robust vision by multi-task learning on monkey visual cortex. Advances in Neural Information Processing Systems 2, 739–751 (2021).

25. I Gusti Bagus, A. M., Marques, T., Sanghavi, S., DiCarlo, J. J. & Schrimpf, M. Primate Inferotemporal Cortex Neurons Generalize Better to Novel Image Distributions Than Analogous Deep Neural Networks Units. (2023).

26. Kingma, D. P. & Welling, M. Auto-Encoding Variational Bayes. Preprint at 10.48550/arXiv.1312.6114 (2022) DOI: 10.48550/arXiv.1312.6114.

27. Srivastava, N., Hinton, G., Krizhevsky, A., Sutskever, I. & Salakhutdinov, R. Dropout: A Simple Way to Prevent Neural Networks from Overfitting. Journal of Machine Learning Research 15, 1929–1958 (2014).

28. Xie, Q., Luong, M.-T., Hovy, E. & Le, Q. V. Self-training with Noisy Student improves ImageNet classification. Preprint at 10.48550/arXiv.1911.04252 (2020) DOI: 10.48550/arXiv.1911.04252.

29. Toosi, T. & Issa, E. B. BRAIN-LIKE REPRESENTATIONAL STRAIGHTENING OF NATURAL MOVIES IN ROBUST FEEDFORWARD NEURAL NETWORKS. in 11th International Conference on Learning Representations, ICLR 2023 vol. 27 (Springer US, 2023). DOI: 10.1038/s41593-024-01758-5.

30. Driscoll, L. N., Shenoy, K. & Sussillo, D. Flexible multitask computation in recurrent networks utilizes shared dynamical motifs. Nat Neurosci 27, 1349–1363 (2024) DOI: 10.1038/s41593-024-01668-6.

31. Bastos, A. M. et al. Canonical Microcircuits for Predictive Coding. Neuron 76, 695–711 (2012) DOI: 10.1016/j.neuron.2012.10.038.

32. Hong, H., Yamins, D. L. K., Majaj, N. J. & DiCarlo, J. J. Explicit information for category-orthogonal object properties increases along the ventral stream. Nat Neurosci 19, 613–622 (2016) DOI: 10.1038/nn.4247.

33. Burgess, C. P. et al. Understanding disentangling in $\beta$-VAE. Preprint at 10.48550/arXiv.1804.03599 (2018) DOI: 10.48550/arXiv.1804.03599.

34. Ye, Z. & Wessel, R. Speed modulations in grid cell information geometry. Preprint at 10.1101/2024.09.18.613797 (2024) DOI: 10.1101/2024.09.18.613797.

35. Kayaert, G., Biederman, I., Beeck, H. P. O. D. & Vogels, R. Tuning for shape dimensions in macaque inferior temporal cortex. 22, 212–224 (2005) DOI: 10.1111/j.1460-9568.2005.04202.x.

36. Higgins, I. et al. Unsupervised deep learning identifies semantic disentanglement in single inferotemporal neurons. 1–14 (2020) doi:10.1038/s41467-021-26751-5 DOI: 10.1038/s41467-021-26751-5.

37. Pedregosa, F. et al. Scikit-learn: Machine Learning in Python. Journal of Machine Learning Research 12, 2825–2830 (2011).

38. Doob, J. L. The Limiting Distributions of Certain Statistics. The Annals of Mathematical Statistics 6, 160–169 (1935) DOI: 10.1214/aoms/1177732594.

39. Bishop, C. M. Pattern Recognition and Machine Learning. (Springer New York, NY, 2006).

40. De, A. G. et al. GPflow: A Gaussian Process Library using TensorFlow. Journal of Machine Learning Research 18, 1–6 (2017).

41. Wilk, M. van der et al. A Framework for Interdomain and Multioutput Gaussian Processes. Preprint at 10.48550/arXiv.2003.01115 (2020) DOI: 10.48550/arXiv.2003.01115.

42. Nejatbakhsh, A., Garon, I. & Williams, A. Estimating noise correlations across continuous conditions with Wishart processes. Advances in Neural Information Processing Systems 36, (2024).

